# PRIEST - Predicting viral mutations with immune escape capability of SARS-CoV-2 using temporal evolutionary information

**DOI:** 10.1101/2023.08.11.552988

**Authors:** Gourab Saha, Shashata Sawmya, Md. Ajwad Akil, Arpita Saha, Sadia Tasnim, Md. Saifur Rahman, M. Sohel Rahman

## Abstract

The dynamic evolution of the SARS-CoV-2 virus is largely driven by mutations in its genetic sequence, culminating in the emergence of variants with increased capability to evade host immune responses. Accurate prediction of such mutations is fundamental in mitigating pandemic spread and developing effective control measures. In this study, we introduce a robust and interpretable deep-learning approach called PRIEST. This innovative model leverages time-series viral sequences to foresee potential viral mutations. Our comprehensive experimental evaluations underscore PRIEST’s proficiency in accurately predicting immune-evading mutations. Our work represents a substantial step forward in the utilization of deep-learning methodologies for anticipatory viral mutation analysis and pandemic response.

## 1 Introduction

The SARS-CoV-2 virus had swept across the world, causing a global pandemic and leading to the unprecedented COVID-19 crisis. While the humanity seems to have survived the worst situation, COVID-19 still impacts global health and society. Viral diseases like COVID-19 are caused by the rapid evolution and virulence of the virus (in this case, SARS-CoV-2). We can follow the evolution and transmission of a virus by identifying and comprehending the genetic changes it undergoes in time. This is crucial, particularly to facilitate vaccine discovery and therapeutics, as we need to develop methods capable to predict future variants that are powerful enough to escape immunity.

With the rise of more sophisticated high throughput sequencing technologies and tremendous progress in machine learning techniques in different fields, various effective computational methods have been developed and proposed to analyze viral evolution and predict viral escape. Salama et al. proposed a neural network-based model to predict the point mutation that takes place by aligning the primary RNA sequence of the Newcastle virus [1]. Yin et al. modeled sequential data of the influenza A virus in the form of time series data and used the LSTM network with attention to predict mutation at any specific residue site [2]. Takwa et al. [3] used a sequence-to-sequence LSTM-RNN model to predict mutations in RNA sequence evolution for successive generations and presented a proof of concept by applying their method on two datasets: Newcastle Disease Virus (NDV) and influenza virus.

Despite being a relatively newer candidate, the progress on this front with respect to SARS-Cov-2 is also fascinating. Bai et al. developed a coarsegrained model and calculated the change of free energy of different single-site or combined-site mutations and predicted potential mutation sites of SARS-CoV-2 [4]. Maher et al. developed a pipeline using different methods, such as, a bidirectional LSTM model, to predict which individual amino acid mutations in SARS-CoV-2 would become more prevalent and contribute to future variants [5]. Zhou et al. used phylogenetic tree-based sampling methods of SARS-CoV-2 viral sequences combined with temporal information and a transformer model to predict mutation at specific sites [6].

Besides mutation prediction, substantial progress is also noticed in the literature in viral escape prediction with probabilistic-based and deep-learning methods. Hie et al. used Bidirectional LSTM models to identify viral escape mutations of influenza hemagglutinin, HIV-1 envelope glycoprotein (HIV Env), and SARS-CoV-2 Spike viral proteins by comparing the mutations of the protein sequences with changes in structure and semantics of the human language [7]. Thadani et al. introduced an interpretable framework incorporating evolutionary data, structural features, and residue dissimilarity properties to predict viral immune escape of SARS-CoV-2 [8].

In this work, we developed a novel interpretable, and explainable deep-learning framework using transformer attention mechanism [9] alongside other techniques in order to predict mutations in the SARS-CoV-2 virus. We validated our predictions by predicting variants ahead of their time. In particular, we corroborated our model’s efficacy by preemptively predicting real-world variant emergence ahead of time. Moreover, our predicted variants were virulent enough to escape immunity which is a great concern and demands further (wet lab) experimental validation. We argue that our method can be applied to predict mutations at specific sites just from sequence data alone, and it performs competitively with existing methods. Thus, the key contributions of this study are twofold:

1. A model with high efficacy in predicting mutations in the SARS-CoV-2 virus that can also be applied in other scenarios, i.e., evolution in other viruses.
2. A methodology that does not restrict itself to known evolutionary paths and can generalize well enough to never before seen cases.

## 2 Materials and Methods

### 2.1 Dataset Preparation

#### 2.1.1 Dataset Collection and Preprocessing

We used sequences of SARS-CoV-2 available in GISAID [10]. We extracted the primary sequences and Pango lineages from their database and collected sequences up to the first quarter of 2022. We divided the sequences into three primary classes based on their variants (Table 1) as follows: (i) benign variants (i.e., the ones whose Pango Lineage did not fall under any of the known variants that were marked deadly at different points in time during the COVID-19 pandemic), (ii) variants being monitored, and (iii) variants of concern.

**Table 1:** Overview of the broad category of Variants used in this study. We consider three variant categories. The names of the variants (if available) are mentioned along with the labels assigned to the categories for the purpose of classification.

| Class | Variants | Label |
| --- | --- | --- |
| Benign Variant | - | 0 |
| Variants being Monitored (VbM) | Alpha, Beta, Delta, Gamma, Epsilon, Eta, Iota, Kappa, Zeta, Mu | 1 |
| Variants of Concern (VoC) | Omicron | 2 |

Our preprocessing steps are inspired by that of Tempel [2], which presents a mutation prediction pipeline for influenza A viruses. However, unlike Tempel, which could leverage 26 years’ of influenza dataset (containing HA proteins of subtypes H1N1, H3N2, and H5N1), we had SARS-CoV-2 protein sequence data for a few years only, i.e., for the period 2019-2022. Therefore, we had to redesign the methodology significantly. We used quarters as our timesteps. So, we have four quarters in a year with Qn representing the *n*th quarter (i.e., Q1: January-March, Q2: April-June, Q3: July-September, Q4: October-December). Breaking down the timesteps of each year into quarters led to a total of 10 timesteps. This adjustment had two major impacts on our study. The quarter-level granularity allowed us to construct longer time series thereby improving the training. Additionally, shorter timesteps allowed us to tackle the challenges of SARS-CoV-2’s rapid mutations [5, 11–14].

The sequences were of varying lengths but generally comprised more than 1000 amino acid residues. Performing multiple sequence alignment on strings with such length and quantity is very slow and computationally expensive (particularly with the resource constraint setting we have to work on). Heuristic methods for MSA are faster but are error-prone [15, 16] and would introduce errors in our input data. Thus, we opted for padding the protein sequences. The length of most of the sequences is below 1280. As such, we marked the ones greater than 1280 as outliers and excluded them from further consideration. All the mutation sites of concern are contained within the first 1228 residues. Hence, sequences that were less than 1230 were also removed to ensure all mutation sites were present in each sequence. We padded out the rest of the sequences (our library) to convert all of them into sequences of uniform length. Thus, we added a maximum of 50 amino acid residues to our sequence (roughly 4% of the total sequence length). This process of padding did not introduce any additional noise or error, as the mutation sites of concern were not modified. Statistics of each timestep are provided in Table 2.

**Table 2:** Number of protein sequences collected per class grouped by years and quarters.

| Timestep | Year | Quarter | Count by Classes |  |  |
| --- | --- | --- | --- | --- | --- |
|  |  |  | Class 0 | Class 1 | Class 2 |
| 1 | 2019 | 4 | 4 | 0 | 0 |
| 2 | 2020 | 1 | 7485 | 32 | 0 |
| 3 | 2020 | 2 | 17654 | 43 | 0 |
| 4 | 2020 | 3 | 21580 | 923 | 0 |
| 5 | 2020 | 4 | 38942 | 8491 | 0 |
| 6 | 2021 | 1 | 67745 | 57452 | 5 |
| 7 | 2021 | 2 | 23758 | 109051 | 1 |
| 8 | 2021 | 3 | 6648 | 254269 | 1 |
| 9 | 2021 | 4 | 4330 | 200097 | 53209 |
| 10 | 2022 | 1 | 625 | 2660 | 38255 |

#### 2.1.2 Time Series Construction

At each timestep, we grouped the protein sequences by the classes of different SARS-CoV-2 variants as detailed in Table 2. Since no evidence suggests that variants mutate into older variants, we assume the same in our study. However, much is still unknown about the exact nature of SARS-CoV-2 evolution. So, we should note that the existence of such cases (mutating into older variants) can be easily handled in PRIEST. Our study aims to understand possible changes in the current variants that may lead to next-generation variants. Moreover, the shift in the percentage of each variant collected over time reinforces our assumption about the virus mutation. Therefore we constructed the time series by linking these protein sequences under special conditions as stipulated below.

Let 𝓁_*i*_ and 𝓁_*i*+1_ be the labels of the protein sequences where 𝓁_*i*_, 𝓁_*i*+1_ ∈ {0, 1, 2}. The groups of sequences with label 𝓁_*i*_ can be linked to the groups of sequences with label 𝓁_*i*+1_ if and only if 𝓁_*i*_ ≤𝓁_*i*+1_. This constructs a graph. From now on, we will refer to this as time series graph.

At each timestep, we randomly sample protein sequences from all possible allowed variants. However, whether we can sample from a variant group depends on which group we sampled from in the previous timestep. For example, if we sample a sequence from Variants Being Monitored (VbMs) at timestep *i*, we can only sample from Variants being Monitored (VbMs) and Variants of Concern (VoCs) at timestep *i* + 1. We can ensure this by following the links in the time series graph. In addition, we performed this sampling with replacement to cover for the lack of enough data of a specific variant at certain timesteps. For instance, at timestep, *i*, the probability of picking a protein sequence of a certain variant is weighted by the number of samples in the allowed group of sequences at any given timestep. This weighted sampling allows us to focus on the dominant variants at each timestep while keeping the others relevant. Eventually, this allows us to capture the variants’ state better.

#### 2.1.3 Input Representation

Now that we have our time series data, we need to convert this into trainable data for our deep-learning models. For any given protein sequence time series 𝒮 and mutation site *m*, we take the three overlapping trigrams *t*_1_, *t*_2_, *t*_3_ containing the mutation site, *m* such that *t*_3_, *t*_2_, and *t*_1_ contain *m* at the first, second and third position respectively (Figure 2). Let us assume we are constructing the time series for site 484, and the protein sequence in Figure 2 is the sequence for timestep 𝒯. The trigrams that contain the site are VEF (in purplr), EGF (in green), and GFN (in yellow). We then take the respective ProtVec [17] representations of the trigrams 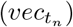. The size (*dim*_*vec*_) of each vector of the trigrams is 100. We aggregate them to get the representation of that specific mutation site at timestep 𝒯. As such, the time series of vector representations at a particular mutation site *m* for a given protein sequence time series 𝒮 that will be input to PRIEST can be represented by 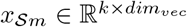 where *k* = the number of time steps and *dim*_*vec*_ = 100 and 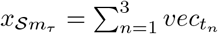.

**Fig. 1:**
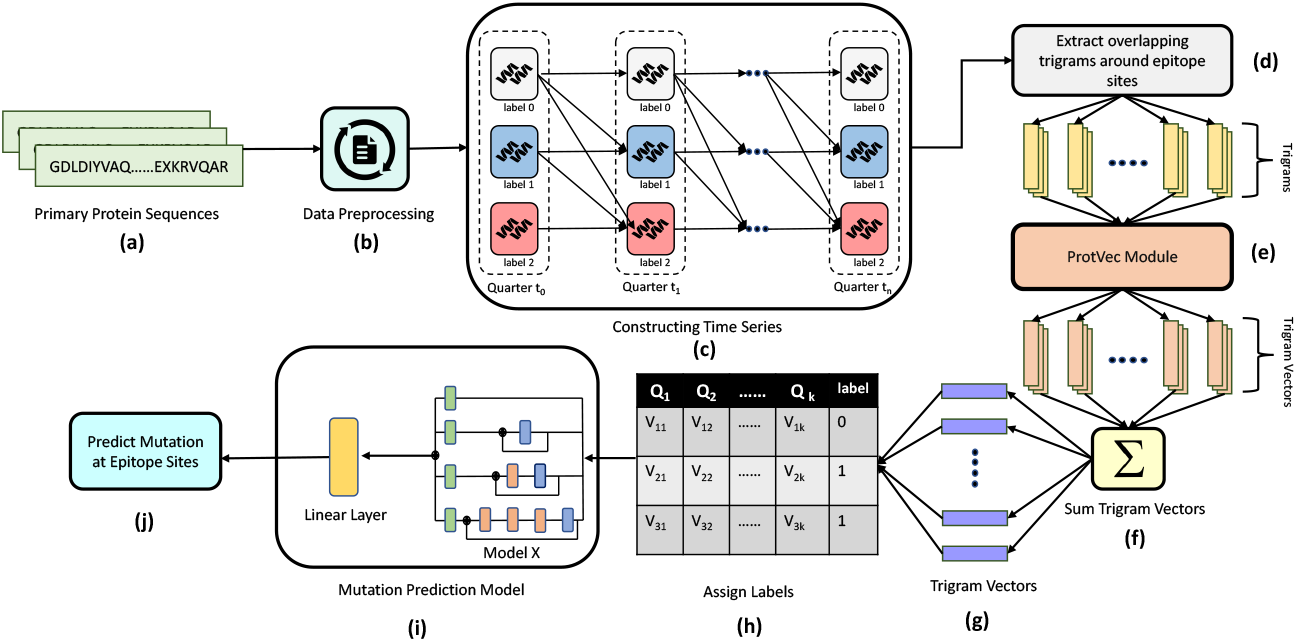
An overview of our Mutation Prediction Pipeline. (a) -(b) indicate protein sequence collection, clean up, and required preprocessing; (c) shows time series construction by modeling the time series in terms of quarters from the fourth quarter of 2021 to the fourth quarter of 2022; (d) -(g) represent steps to create training data for PRIEST, we transformed the training sequences into embeddings that can be used for training. In (h) we assigned the labels to our training data; (i) indicates training PRIEST and (j) represents the mutation prediction at respective epitope sites.

**Fig. 2:**
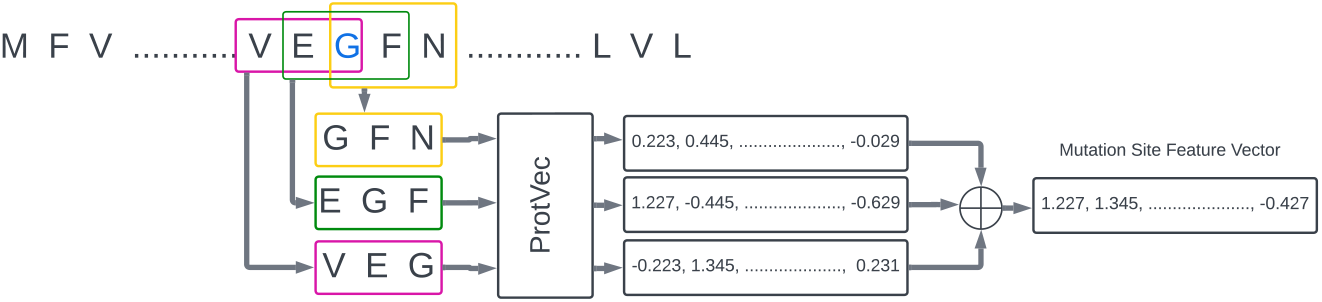
Extraction of feature representation of any given mutation site. We use a sliding window to extract three trigrams around that position. In the sequence above, to get the feature representation of the site marked with blue color, we extract VEG (in purple), EGF (in green) and GFN (in yellow). We use ProtVec to extract feature representation of each of the three trigrams. We aggreagte them to obtain the feature represnetation of the position.

**Fig. 3:**
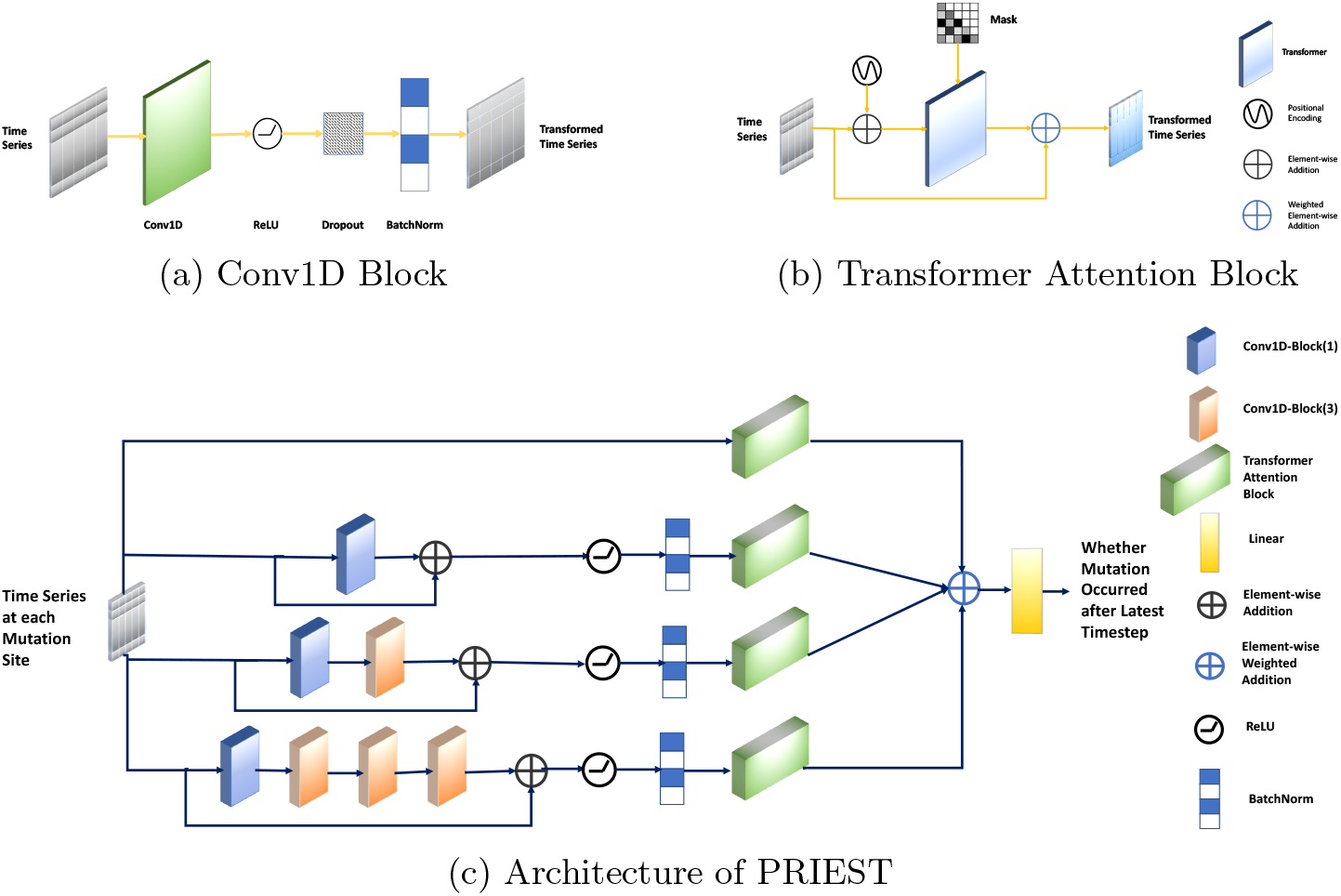
PRIEST Architecture with Detailed View of All the Components. (a) The architecture of 1D convolution block used for feature transformation. (b) The architecture of the transformer attention block. (c) A schematic diagram of the overall architecture of PRIEST.

#### 2.1.4 Label Assigning

To train PRIEST, we need to assign a label to each data point. If there is a difference of residue at site *m* between protein sequences at the final and the penultimate time steps for a protein sequence time series 𝒮, we consider this as a mutation and assign a label 1 to the corresponding time series of vector representations. Otherwise, we assign a label 0. We construct the dataset with time steps 3, 6 and 9. The count of positive and negative data points for training, validation, and testing set for time steps 3, 6 and 9 are reported in Table : 3.

### 2.2 Overview of PRIEST

#### 2.2.1 The Conv1D Block

One of the key components of PRIEST is the Conv1D block. This block takes a time series at mutation site 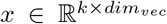 as input and passes it through a 1D convolution layer [18] of kernel size *k* and padding *p* as seen in Equation 1. This is followed by a ReLU [19] activation function and a subsequent dropout [20] layer. Additionally, we applied batch normalization [21] to prevent internal covariance shift. We obtain 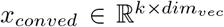 as output. These transformations can be represented by (the 3 equations presented in) Equation 2.

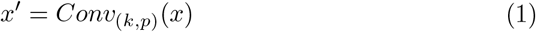

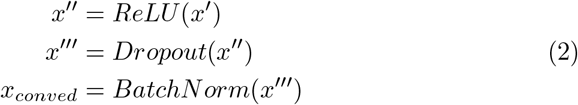

#### 2.2.2 The Transformer Attention Block

The second key component is the transformer attention block. This is close to the Self-Attention module introduced in [22]. Positional encoding [9] is applied to our input to introduce relative and absolute positional information about the timesteps and can be represented using the Equations 3 and 4.

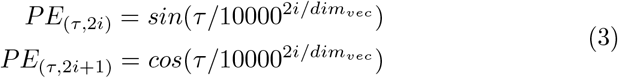

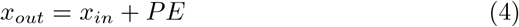

The transformed input 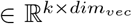 is passed through a transformer encoder [9] with *l* layers and *h* attention heads. We used an attention mask that prevents the timesteps of the input time series from attending to future timesteps. The rationale behind the mask is that evolution is dependent on the environment and thus has plenty of variations. As such, we do not want PRIEST to make decisions about past timesteps using the information of future ones. The output is represented using 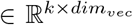. We multiply the output of the transformer encoder by *α* and the original input by 1 −*α* to obtain the output. Here, *α* is a learnable parameter.

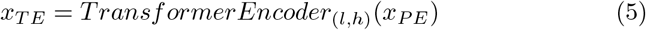

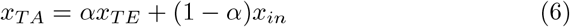

#### 2.2.3 PRIEST Architecture

Now we are ready to discuss the overall architecture of PRIEST. PRIEST incorporates the ideas of the Inception module introduced in [23] and the modifications introduced in [24]. There are 4 channels. The first channel applies no convolution operations. The second channel applies one convolution operation with a Conv1D Block of kernel size 1. In the third channel, we apply two Conv1D Block operations, one of kernel size 1 and the second of kernel size 3. Finally, the fourth channel applies four consecutive Conv1D Block operations with kernel sizes 1, 3, 3, and 3 respectively. We add the input *x*_*in*_ to the output of each of the convolution channels via a residual connection [25]. We apply ReLU and Batch Normalization after the residual connections. We apply the transformer attention mechanism described previously to each of the channels. We represent the outputs by 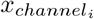 where *i* ∈ [1, 4]. We multiply them by weights *w*_*i*_ and add them to obtain *x*_*out*_ (Equation 7).

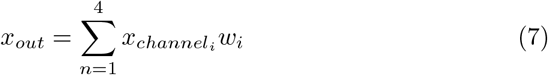

We extracted the most recent timesteps of *x*_*out*_ and pass them through a linear layer to calculate the predicted score, ŷ. We calculated the cross-entropy loss between ground truth *y* and ŷ and this can be represented by Equation 8.

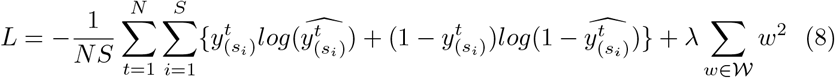

Here, *N* is the number of training examples and *S* is the number of mutation sites. Additionally, *s*_*i*_ represents *i*-th mutation site. The second term in the loss function is the L2-regularization term. In this term, *λ* is the regularization parameter and it is a hyperparameter, i.e., it is not a learnable parameter. On the other hand, 𝒲 represents the set of learnable parameters.

### 2.3 Mutation Quality Testing

In order to validate the effectiveness of PRIEST, we also generated new mutated sequences for Q2 2022 by mutating sequences of Q1 2022. We selected the mutation site following [26, 27]. Then we used PRIEST to predict mutation at those specific sites for Q1 2022 sequences. In any given mutation site, often more than one residue can replace the existing one. We performed this mutation with equal probability among the possible replacement residues. To understand the strength and broad classification of the newly mutated sequences, we classified the newly mutated sequences using the classification pipeline we developed to classify the three variants from sequence data -Unknown Variants, Variants being Monitored, and Variants of Concern. Details related to the classification experiment can be found in section 7.1 of the supplementary part of the paper. The Sequences that were classified as Variants of Concern (VoCs) are used for further analysis.

To assess the quality of the generated sequences, we performed four experiments. These experiments are briefed in the following sections.

### 2.3.1 Calculating Sequence Embedding Distance

We calculated the mean of the ProT5 vector representations of the actual Q2 sequences. We represent this by *mean*_*true*_. We also find *mean*_*gen*_, the mean of the generated sequences. Then we used Euclidean Distance and Cosine Dissimilarity to calculate the distance between the two types of embeddings.

#### 2.3.2 Clustering and Pairwise Distance Computation

While it is useful to find the overall distance between the sequences, the sequences themselves do have some variations among themselves for a particular variant. To capture these differences and variations, we ran clustering algorithms. We then computed the pairwise distance between the clusters to capture the details of the differences even further.

For clustering, we first ran PCA on the actual existing sequences to find the principal components. We measured the variance caused by each of the components. The first two components had the highest degree of variance. We took these components and divided the sequences into clusters using Gaussian Mixture Models (GMM). We chose GMM over other methods because it allows us to determine the clusters and does not need to perform any additional tests to find the distribution of the clusters. GMM uses the Bayesian Information Criterion (BIC) score to identify the number of clusters. In this case, the lower BIC score is better. Another important hyperparameter of this model is the covariance type, which controls the degrees of freedom to define the shape of each cluster. There are four covariance types: spherical, tied, diagonal, and full. To obtain the desired number of clusters and the covariance type, we first used grid search from the Scikit-learn package, considering up to nine components and all covariance types. We then generated the BIC score for each covariance type with respect to all the clusters. We observed from the plots of the BIC score vs. the number of clusters that the score for the diagonal type of covariance is the lowest, so this is our desired covariance type. To determine the number of clusters, we calculated the gradients of the BIC score. We observed that the gradient for the diagonal type of covariance did not improve after seven components. Thus, we selected our desired number of components to be seven for performing the clustering. Further details can be found in the supplementary section 7.3.2.

We trained a GMM model with the values identified after hyperparameter tuning (BIC and covariance type analysis). We used the principal components of the original sequences for training purposes. Then we fitted our generated sequence’s principal components into the clusters. For a quantitative comparison of the clusters, we used the NearestCentroid algorithm [28]. to fit the original sequences and the generated sequences with their respective cluster labels. Then we compared the centroids considering both Euclidean distance and cosine similarity.

#### 2.3.3 Calculating KL Divergence Score

To calculate KL Divergence Score, we selected the existing sequence (Q2 2022) embeddings and our generated embeddings. We grouped them by the common clusters in the existing and generated sequence clusters. We then split both the embeddings data in a 70:30 ratio for the training and testing part to fit a Gaussian distribution to our clusters. We used Gaussian distribution as we used GMM to cluster the embeddings. We used KernelDensity estimation of Scikit-learn to fit Gaussian distributions using the training samples. Then we obtained the log of probability density values using the testing samples and the probability density values of both types of sequence embeddings. After that, we computed the forward KL Divergence between the two distributions. We considered the distribution of original sequence embeddings, *C*_*e*_, as our reference distribution and the one for the generated embeddings *C*_*g*_ as our approximation. The full algorithm for computing the divergence score is given in Algorithm 1.

##### Algorithm 1 KL-Divergence on cluster distribution

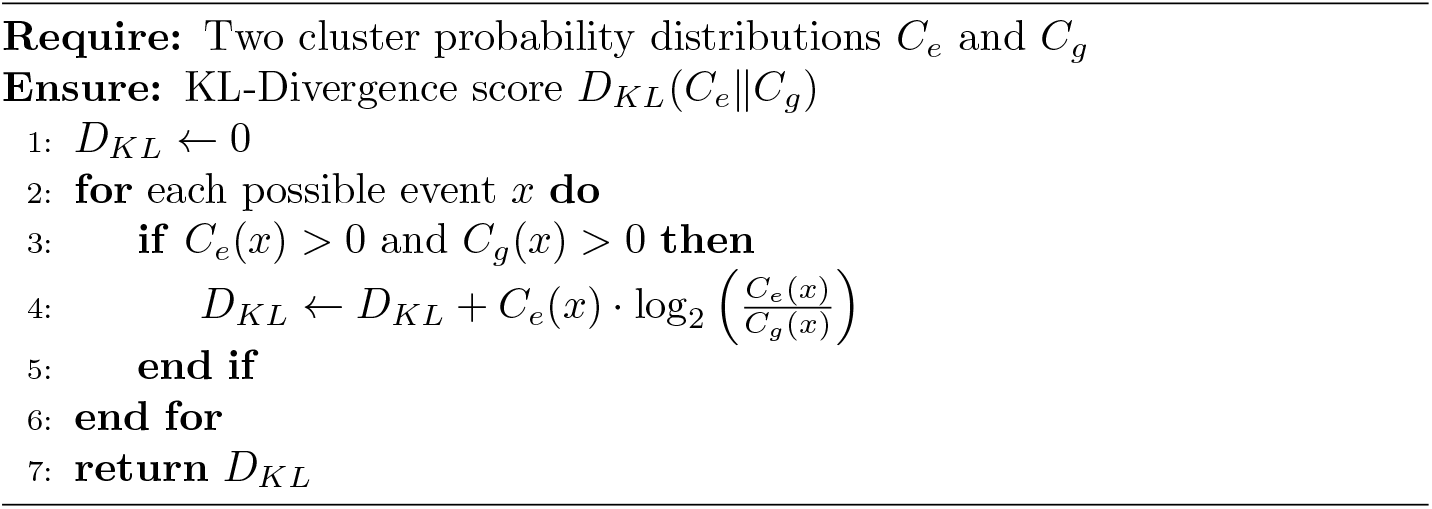

#### 2.3.4 Calculating Immunescape Score

We also experimented with models that can quantify how much a mutated sequence is likely to escape the immune system of the host. We used the EVEScape index introduced in [8], which incorporates fitness predictions from evolutionary models and features related to structural information to determine antibody binding to the mutated proteins, as well as the distance between normal and mutated residues. It uses a probabilistic model that generates a log-likelihood score, giving us the probability of a single amino acid substitution leading to immune or antibody escape. Three separate (likelihood) scores are calculated first as follows[8], and the final score is calculated as the product of these three. The first score gives the likelihood of maintaining the mutation’s fitness by training a deep generative model. The second score gives the likelihood of the mutation being accessible to the antibody. And the third score gives the likelihood of the mutation disrupting binding to the host antibody.

Aside from the original sequences and the generated mutated sequences (generated by PRIEST as mentioned in Section 2.3), we also generated random mutations by modifying the original sequences as follows. With some fixed probability *p*, we selected whether a mutation would occur at a given mutation site. In case of more than one possible candidate, we performed this mutation with equal probability among the possible replacement residues. Finally, we calculated respective Immunescape scores for both the random mutations and our generated mutations using the aforementioned EVEScape model and compared them.

### 2.4 Evaluation Metrics

In this study, we perform our evaluation from two different angles. First, we evaluate our model, PRIEST to compare it with other models from machine learning point of view. In parallel, we also want to examine the quality of mutation of the mutated sequences generated by PRIEST. So, we use (i) Model Evaluation Metrics and (ii) Mutation Quality Testing Metrics as described below.

#### 2.4.1 Model Evaluation Metrics

For evaluating PRIEST and other baseline models, we used accuracy, F-score, Matthews Correlation Coefficient (MCC), Receiver Operating Characteristics (ROC) curve, and Precision-Recall(PR) curve. For the ROC and PR curves, we consider the area under these curves. These are also known as AUROC and AUPRC, respectively. We consider MCC the primary metric of concern as it provides a better measure of the performance of a model for binary classification over accuracy and F-score despite data imbalance [29]. From Table 3, we can see a heavy imbalance in our dataset for train, validation, and testing sets for all values of *k*. Hence, MCC is a good performance metric to assess PRIEST’s performance. We also use F-score as our secondary metric for this particular concern.

**Table 3:** Details of train, validation, and testing set for time steps (*k*) 3, 6 and 9.

| Data Split | k=3 |  | k=6 |  | k=9 |  |
| --- | --- | --- | --- | --- | --- | --- |
|  | 0 | 1 | 0 | 1 | 0 | 1 |
| Train | 128631 | 318369 | 127083 | 319917 | 125311 | 321689 |
| Validation | 7292 | 22508 | 8186 | 21614 | 9579 | 20221 |
| Test | 7483 | 22317 | 8556 | 21244 | 8518 | 21282 |

#### 2.4.2 Mutation Quality Testing Metrics

As detailed throughout Section 2.3, we performed additional tests to verify the quality of mutations generated by PRIEST. For calculating the difference between the generated and existing sequences in Q2 2022, we used Euclidean distance and Cosine dissimilarity as our metrics. They allow us to find the difference between the representations of two entities in a given vector space. Additionally, we used the KL-Divergence score to find the difference between the distributions of clusters of similar types of sequences in existing and generated sequences. Finally, we used the EVEScape score to assess the chance of our generated sequences to evaluate their potential threats.

### 2.5 Implementation and Training

We used CNN [30], RNN [31], LSTM [32], GRU [33], Tempel (both in temporal attention and dual-attention form) and Tempo [6] as our baselines. We used Pytorch and Scikit-learn for implementing our architectures and evaluating metrics respectively.

The CNN architecture comprised 1D convolution layers followed by 1D max pooling layers. We used 3 windows of sizes 1, 2, and 3 for convolution and a dropout rate of 0.2. For the remaining baselines, a hidden size of 128 and a dropout rate of 0.5 were used. We used Tempo with two transformer encoder layers and five attention heads.

In PRIEST, we used 1D Convolution Layers of primarily 2 types-one with kernel size 1 and padding size 0, and the other with kernel size 3 and padding size 1. For our transformer encoders, we used 2 transformer encoder layers and 5 attention heads. We applied a dropout of 0.5 to all layers and 0.1 in positional encoding.

For maintaining consistency in performance measurement, we trained all the models for 50 epochs with a batch size of 256. We used Adam [34] as our optimizer with learning rate 1 *×* 10^−3^. To prevent overfitting and ensure an additional boost to our performance, we used a cyclic learning rate scheduler [16].

### 2.6 Code, environment, and availability

All the training was done on a machine with Intel® Core™ i7-9700F CPU @ 3.00GHz processor with 8 cores and 16GB of RAM, and GeForce RTX 2070 SUPER GPU with 8GB VRAM and Ubuntu 22.04.2. PRIEST is available as an open-source implementation in https://github.com/ComeBackCity/PRIEST.

## 3 Results and Discussion

### 3.1 Efficacy of Mutation Prediction

The performance of all the models used in our experiments is reported in Table 4. As we can see PRIEST has higher MCC than other baseline models for k ∈ {3, 6}. Additionally, for *k* = 9, it still has comparable performance. Moreover, with respect to the other two metrics of concern, PRIEST still outperforms or compares well to other models across all the different time series sequence lengths.

**Table 4:**
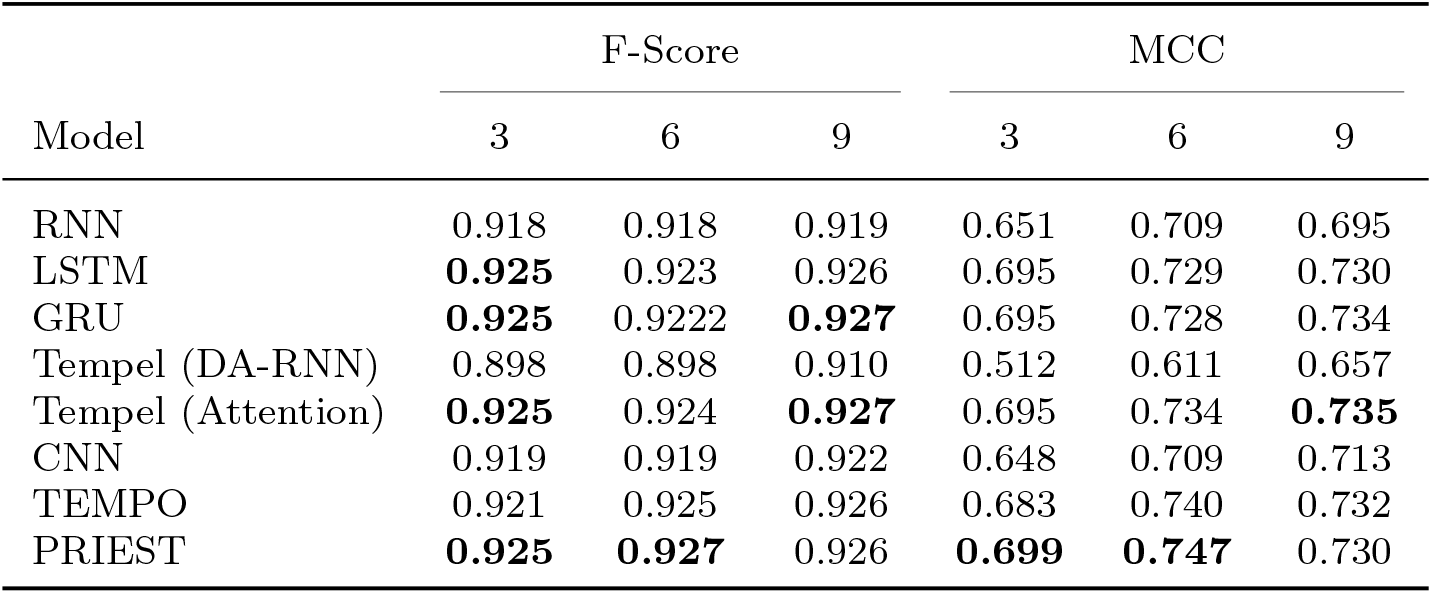
A comparison of evaluation metrics (F-Score and MCC) obtained by PRIEST and other benchmark models on independent test set.

| Model | F-Score |  |  | MCC |  |  |
| --- | --- | --- | --- | --- | --- | --- |
|  | 3 | 6 | 9 | 3 | 6 | 9 |
| RNN | 0.918 | 0.918 | 0.919 | 0.651 | 0.709 | 0.695 |
| LSTM | **0.925** | 0.923 | 0.926 | 0.695 | 0.729 | 0.730 |
| GRU | **0.925** | 0.9222 | **0.927** | 0.695 | 0.728 | 0.734 |
| Tempel (DA-RNN) | 0.898 | 0.898 | 0.910 | 0.512 | 0.611 | 0.657 |
| Tempel (Attention) | **0.925** | 0.924 | **0.927** | 0.695 | 0.734 | **0.735** |
| CNN | 0.919 | 0.919 | 0.922 | 0.648 | 0.709 | 0.713 |
| TEMPO | 0.921 | 0.925 | 0.926 | 0.683 | 0.740 | 0.732 |
| PRIEST | **0.925** | **0.927** | 0.926 | **0.699** | **0.747** | 0.730 |

Tempel uses LSTM as their hidden units. While LSTMs outperform GRUs and RNNs in capturing long-term dependency, they are still outperformed by transformers to capture sequential/temporal information. As these units are sequential, a certain loss of information is expected when we use RNN and similar recurrence-based models. Transformers use self-attention; instead of looking at the timesteps in a recurrent manner, it looks at the whole time series. As such, it is generally more potent in capturing long-term dependencies found in the dataset. While Tempo only leveraged this power of transformers and the attention mechanism that comes with it, PRIEST leverages the power of 1D convolution layers to extract additional features. It uses trainable weight parameters to focus on the extracted features or original time series as required.

The performance of PRIEST deteriorates when *k* = 9. Tempel outperforms PRIEST in that specific case albeit marginally. Here, loss of information due to the inability to capture long-term dependency is actually helpful for Tempel. The longer series might add additional information that is not quite as useful for predictions. On the other hand, Temple might have lost the information about the earlier timesteps and only used the useful ones. Ultimately, the model’s complexity, while benefiting PRIEST in most cases, actually hurts in this particular one.

In terms of AUROC and AUPRC, PRIEST outperforms all other baseline models. This is evident in Figure 4 where We plotted the precision-recall curves and receiver operating characteristic curves; PRIEST has a higher area under both the curves in all cases k ∈ {3, 6, 9} (detailed in supplementary section-7.2.1. Since PRIEST at *k* = 6 outperforms all other models in all scenarios, we will use this for further analysis for the remainder of this study.

**Fig. 4:**
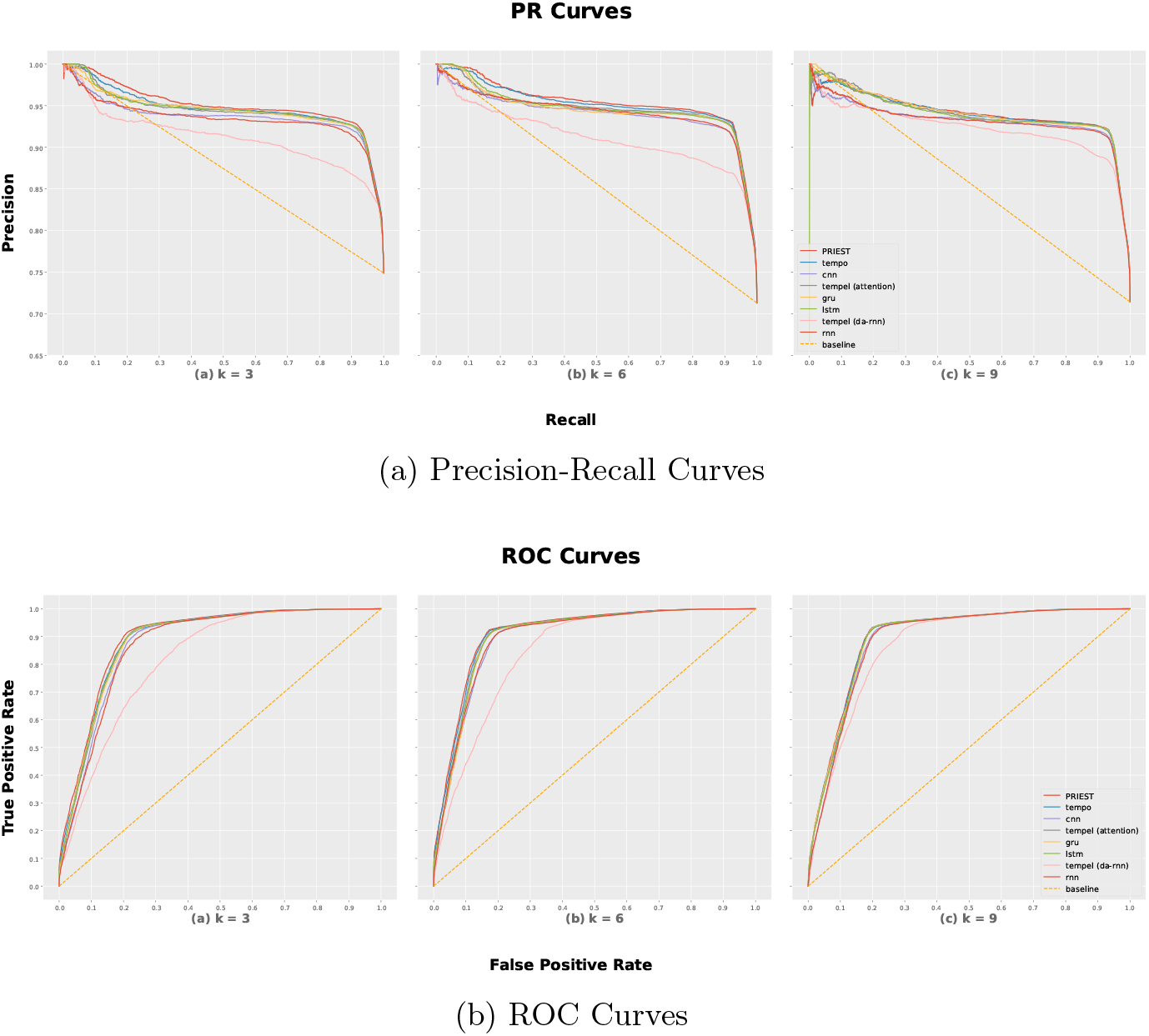
We compute the precision-recall and receiver-operating characteristic (ROC) curve for PRIEST and other baseline models for timestep k∈ {3, 6, 9 }. PRIEST comfortably outperforms the other models by having both greater AUPR and AUROC, respectively.

### 3.2 PRIEST’s Efficacy in Creating Mutated Sequences

To test PRIEST’s ability to detect mutation, we tested it on an independent test dataset to predict the mutations that would arise in Q2 2022. We tested the quality of the mutations that were predicted by PRIEST, we ran the tests as described in Section 2.3. In this section, we are going to describe those results.

#### Embedding Similarity

The Euclidean distance between *mean*_*true*_ and *mean*_*gen*_ was 0.368, while the cosine dissimilarity was 0.024. The results of this study indicate a high level of similarity between the Q2 2022 sequences and the sequences generated in this study, as demonstrated by two different measures.

A lower value of Euclidean distance indicates higher quality as it indicates the closeness of the two sets. The Euclidean distance between the mean values of the two sets of sequences was found to be 0.368, indicating that the two sets of sequences are relatively close to each other in the latent vector space.

Additionally, cosine dissimilarity can vary between 0 (completely similar) to 1 (completely dissimilar). The cosine dissimilarity between the two sets of sequences was measured to be 0.024, indicating a high degree of alignment between the two sets of sequences.

#### Clustering and Pairwise Distance Computation

Our experiments reveal that the generated sequence clusters are highly comparable to the original ones, as seen in Figures 5a and 5b. Most of the clusters present in the set of existing sequences were also observed in the generated set, except for Cluster 4 and Cluster 6.

**Fig. 5:**
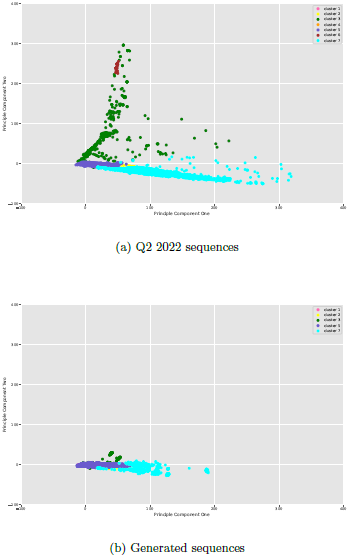
Cluster comparison of actual and generated sequences. Exact coloring is maintained for sequences belonging to the same clusters in both cases.

**Fig. 6:**
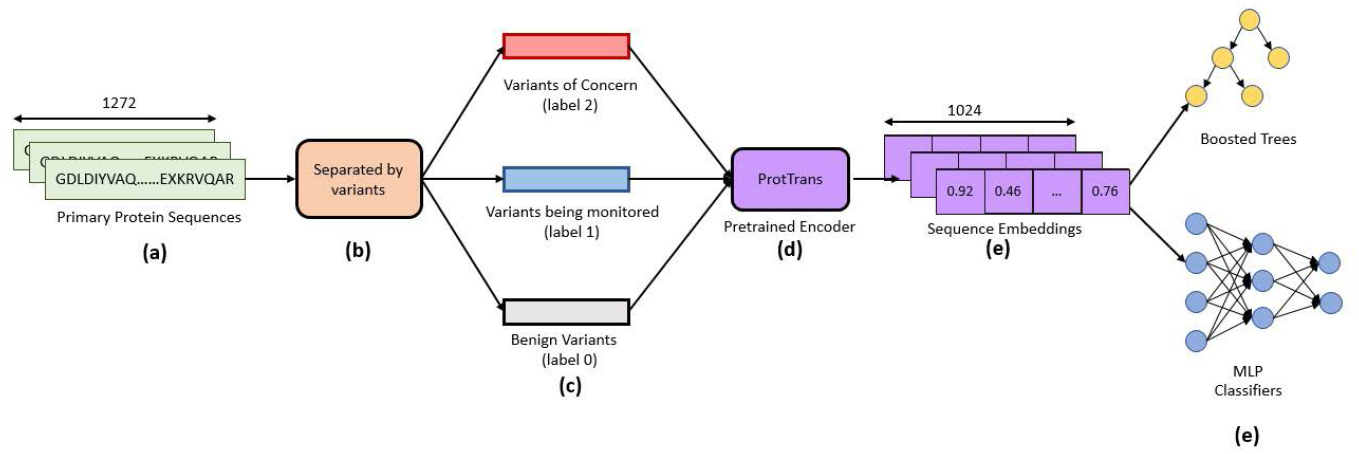
Classification Pipeline

**Fig. 7:**
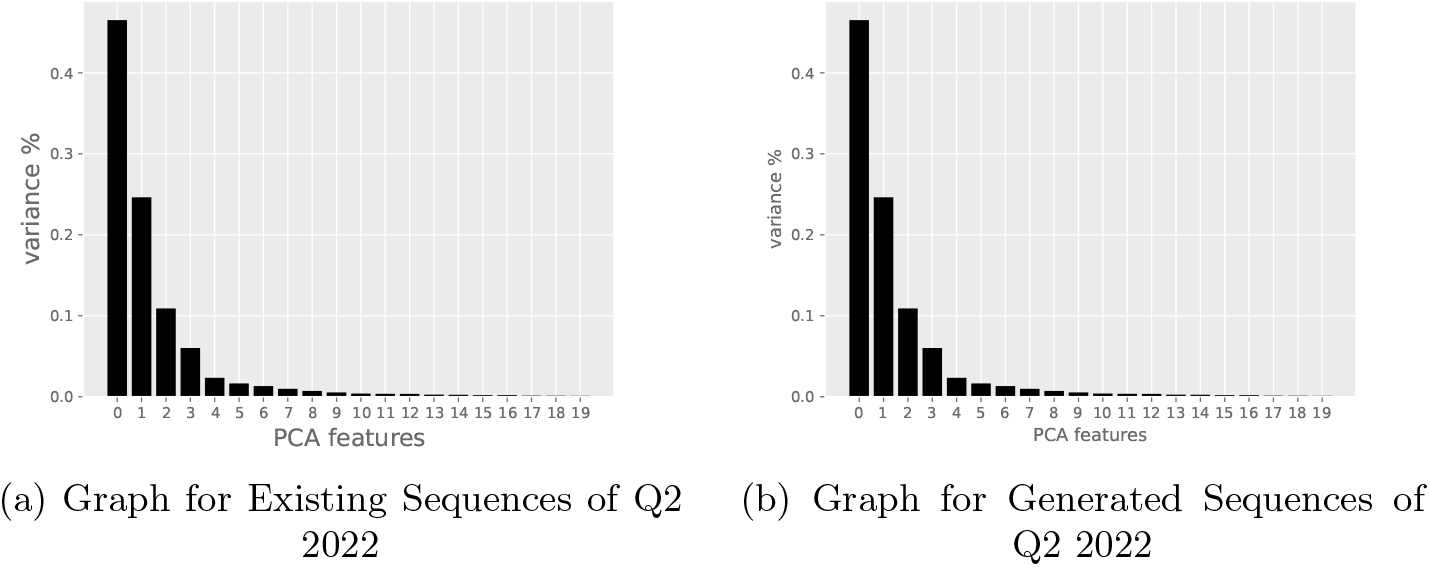
Graphs for GMM Parameter Selection

**Fig. 8:**
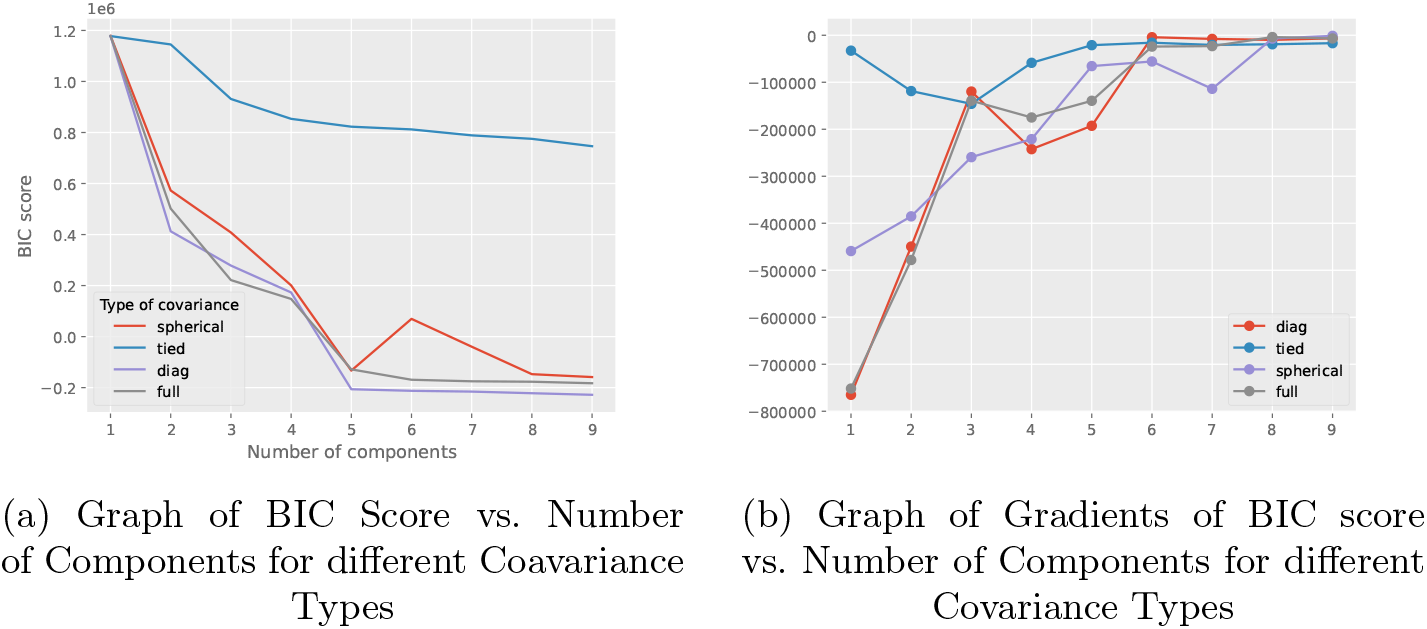
Graphs for GMM Parameter Selection

To further investigate the similarity between the two sets of sequence clusters, Table 11a and Table 11b were used to compare the centroids of each cluster of the actual and generated sequences. Table 11a shows that, for the most part, the centroids of all the generated sequence clusters were closest to their corresponding existing sequence cluster centroids, with the exception of Cluster 3. In fact, for Cluster 3 of the generated sequence, we found that its centroid was closest to that of Cluster 5 of the actual sequence. Likewise, Table 11b reveals that the centroids of generated sequence clusters 1, 2, 5, and 7 were closest to their corresponding actual sequence cluster centroids. Interestingly, Cluster 3 of the generated sequence was again found to be closest to the centroid of Cluster 5 of the actual sequence.

Overall, these findings suggest that the generated and original sequence clusters are indeed close to each other, with only a few anomalies. This anomaly can be attributed to the number of sequences in Cluster 7 of the generated sequences.

#### KL-Divergence Score

To assess the similarity between the distributions of the clusters of the existing and generated sequences, we calculated the KL-Divergence score. The results, presented in Table 5, show that the scores for clusters 1, 4, 5, and 6 are very low, ranging from 0.001 to 0.061. This suggests that the distributions of the clusters of existing sequences and those of the generated ones are quite similar. However, for cluster 7, the KL-Divergence score is 0.292, which is higher than the other clusters. This finding is consistent with the observation made in the analysis of the cluster centroids’ distance, as mentioned earlier. These results provide additional evidence that the generated and original sequence clusters are comparable to each other, with only a few exceptions.

**Table 5:** KL Divergence Score of clusters.

| Cluster Name | KL Divergence Score |
| --- | --- |
| Cluster 1 | $4.897 \times 10^{-5}$ |
| Cluster 2 | 0.049 |
| Cluster 3 | 0.252 |
| Cluster 5 | 0.027 |
| Cluster 7 | 0.177 |

Overall, the KL-Divergence scores indicate that the generated sequence embeddings’ underlying distributions are very close to those of the original sequence embeddings, which is an important finding for applications that require accurate comparisons of sequence data.

#### Immune Escape

We also experimented with models that can quantify how much a mutated sequence is likely to escape the immune system of the host. We used the EVEScape index introduced in [8], which incorporates fitness predictions from evolutionary models, and features related to structural information to determine antibody binding to the mutated proteins, as well as the distance between normal and mutated residues. It uses a probabilistic model that generates a log-likelihood score, giving us the probability of a single amino acid substitution leading to immune or antibody escape. The product of three separate probabilities calculates this score. The first score gives the likelihood of maintaining the mutation’s fitness by training a deep generative model. The second score gives the likelihood of the mutation being accessible to the antibody. And the third score gives the likelihood of the mutation disrupting binding to the host antibody. All the likelihoods are multiplied to get the final score.

Aside from the original sequences and the generated mutated sequences, we also generated random mutations by modifying the original sequences. With some fixed probability *p*, we selected whether a mutation would occur at our selected mutation sites. If the mutation occurs, then we defined the mutation would be a random mutation with equal probabilities for all possible replacements of the residues.

Next, we calculated the Immunescape score for both the random mutations and our generated mutations using the EVEScape model. The details of the calculation can be found in the supplementary section **??**. The scores for each probability of random mutations are shown in Table 6. And for our generated mutations from the proposed model, we obtained the Immunescpae score of − 51.049. Our score is higher than any random mutations, indicating that the mutations generated by using PRIEST may escape the host antibody than any random mutations.

**Table 6:** Immunescape scores against various random mutation probabilities.

| Probability of<br>Random Mutations | Immunescape Score of<br>Random Mutations |
| --- | --- |
| 0.5 | -133.861 |
| 0.6 | -160.697 |
| 0.7 | -187.502 |
| 0.8 | -214.231 |
| 0.9 | -240.987 |

## 4 High Probability Mutation Sites

For this analysis, we took the top 20 sites with the highest mutation probability as predicted by our model. We can see from Table 7 that our model accurately predicts the high-probability mutation sites. Some of the currently known mutations are also shown in the table. While mutations that cause immunescape occur in a specific site, the neutralizing antibody(nAb) that causes this immunescape in the first place also interacts with other sites in the vicinity. Thus, those adjacent sites are also prone to mutations [35]. Our model accurately predicts these as potential mutation sites as well. For example, from Table 7, we can see that K417N is a well-known mutation in site 417. However, our model predicts sites 413-416 as potential high-risk mutation sites. Some of these interactions which are well-known are D414, D415, and F416 [36]. Similar trends can be seen in sites 444 and 445 with well-known interactions being K444 and V445 and the prominent mutation being seen in site 446, G446S. Our mutations are covered within the RBD region of S-glycoprotein. This is the most well-known mutation region in Omicron Variants i.e. the VOCs. The alignment of existing evidence with the predictions made by PRIEST gives validity to its predictions. As such, a site that has been predicted by PRIEST as a high-probability mutation site but has no known mutation associated with it is worth keeping an eye on as these sites are quite likely to undergo mutations as new lineages/variants emerge.

**Table 7:**
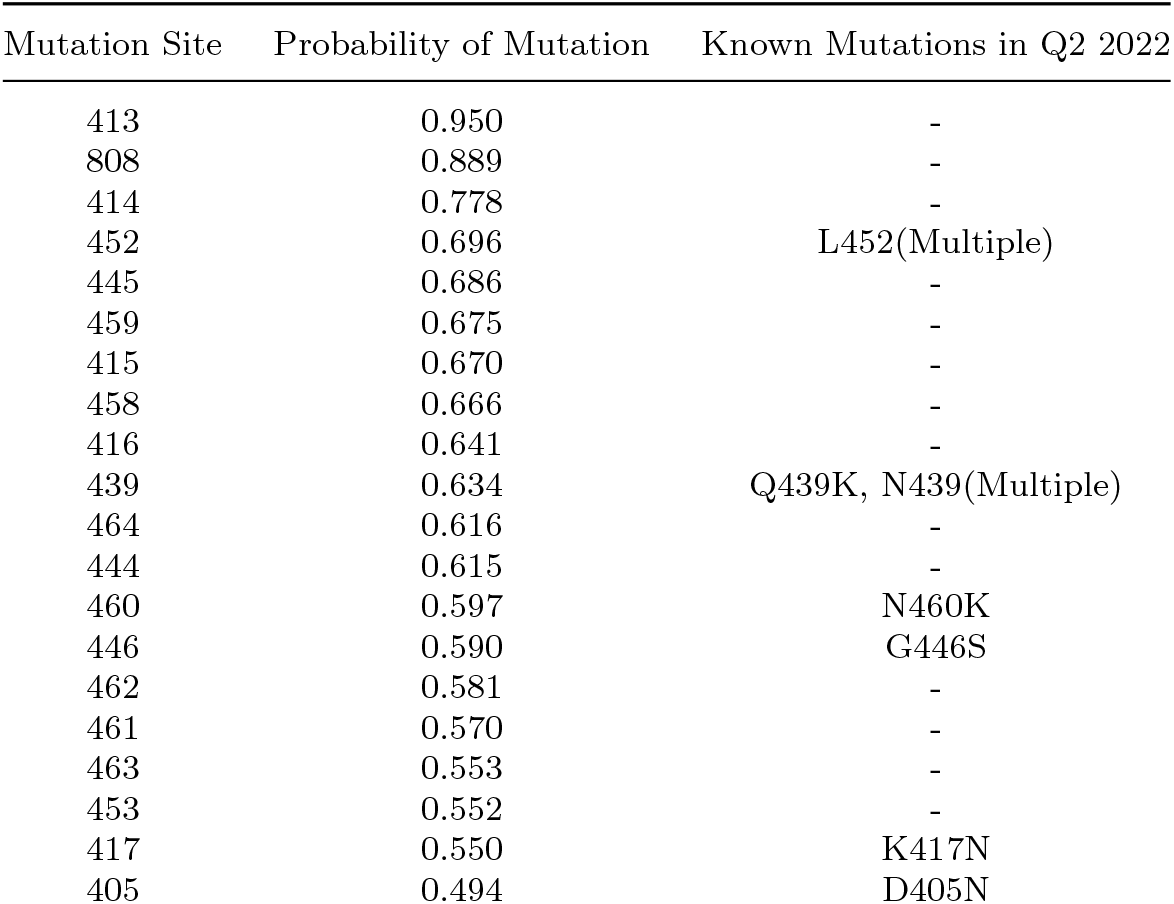
Top 20 mutation sites with the respective probability of mutation as calculated by PRIEST.

**Table 8:** Classes.

| Class | Variants | Label |
| --- | --- | --- |
| Unknown Variant<sup>1</sup> | - | 0 |
| Variants being Monitored | Alpha, Beta, Delta, Gamma, Epsilon, Eta, Iota, Kappa, Zeta, Mu | 1 |
| Variants of Concern | Omicron | 2 |

<sup>1</sup>Benign variants are the ones that didn't have Pango Lineage information.

## 5 Interpretation

During inference, we extracted the attention scores for all the sequences. Additionally, we also extracted the weight each channel received while making the final prediction. We aggregated the attention scores in a weighted manner to calculate the final attention score. Finally, we extracted the top 7 sites from each timestep ranked by the attention score they received.

In Q4 2021, 4 of the 7 sites were located around the RBD region specifically in and/or around sites 452, 453, and 484. These were major mutations found in S-glycoproteins of VOCs [37, 38]. Two of the sites were in the NTD region and the other significant mutation region of S-glycoprotein. A similar trend can be observed, in Q2 2021 and Q3 2021. However, the mutation sites were more evenly distributed between RBD and NTD regions. In timesteps older than Q2 2021, the model focused on sites belonging to primarily the NTD region. The ones which were not part of the NTD region belonged to NSP1 and ORF3. These are 2 of the major regions outside the S-glycoproten mutations.

We can conclude from this discussion that during the prediction of mutation, PRIEST pays attention to the mutation sites belonging to the most vital regions in each timestep. This was achieved without ever passing this information explicitly.

## 6 Conclusion

Mutation prediction in genome sequences is a challenging task at the core of biological research. Moreover, predicting the viral evolution of a dynamic virus like SARS-CoV-2 requires sophisticated methods which can tread in the shifting constraint of immunity [8]. In this study, we presented a novel state-of-the-art method to predict possible mutations in SARS-CoV-2 using historical mutation information. Our method outperforms the existing methods currently used in this field of study. This allows us to deal with the prediction of viral evolution with greater accuracy. Additionally, we present some unique mathematical studies that allow us to test the quality of the new generation of viruses. Not only can these methods be used for future directions of this study, but will also allow us to test relationships among variants of any virus.

Preparation for an upcoming pandemic is impossible if we are not equipped with an early detection mechanism and tools to predict and simulate the future variants of a quickly evolving “germ” of the pandemic. Furthermore, a robust method should likely detect key escape mutations on the shifting grounds of viral immune escape. PRIEST is a flexible method that incorporates evolutionary information to generate such predictions. This allows us to keep up with this continuously evolving virus and allows doctors and researchers to create vaccines as soon as a new variant has arrived. We would no longer need a new variant to be available in nature to create a vaccine for it. This will allow us to save countless lives, something we could not do when COVID-19 originally broke out. Additionally, the simplifying nature of the assumptions we made allows us to use the method for other viruses and pandemics in general.

PRIEST is a general framework for viral mutation prediction. However, it is not without some limitations. We did not evaluate our predictions in the light of Polygenic Risk Scoring (PRS) to associate the disease at the individual level. Second, PRIEST requires sufficient evolutionary information regarding the virus sequence to generate reasonable predictions. Future work will aim at mitigating these limitations and presenting PRIEST as the all-purpose prediction and analysis framework for pandemic preparedness.

## 7 Supplementary information

### 7.1 Classification

#### 7.1.1 Data Collection and Preprocessing

We used SARS-CoV-2 data which are available at GISAID. We extracted the primary sequences and Pango lineages as those are the required information for our tasks. We collected sequences up to the first quarter of 2022. We divided the sequences into three primary classes based on their variants. These are (i) benign variants, (ii) variants being monitored, and (iii) variants of concern. Table 1 shows the classes along with their variants.

We transformed each protein sequence into its hidden representations or embeddings. We used a pre-trained ProT5 model for this transformation. Each representation is a vector V = (*V*_0_, *V*_1_,, *V*_1023_) which has a dimension of 1023.

#### 7.1.2 Classification Model

We used a simple multilayer perception (MLP) network for our purpose. PRIEST consists of 6 hidden layers. The first five hidden layers had 512, 256, 64, 32, and 8 activation units, respectively. Each of these layers had a layer of ReLU as non-linearity after it. The final layer had three activation units, each for one of the classes. A Softmax layer follows this. The first five layers follow the following equation:

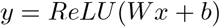

and the final layer follows the following equation:

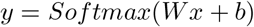

#### 7.1.3 Hardware

All models associated with classification were trained in Google Colab Pro in High RAM Settings. This setting provided us with 24GB of RAM and an instance of Tesla T4 GPU with 16GB VRAM. Models were trained on GPUs for faster training.

#### 7.1.4 Implementation and Training

We transformed the protein sequences into embeddings using ProT5. We divided the data into three splits of 80-10-10 with constant seed for consistency of testing across the models. Apart from conversion to embeddings, all other preprocessing steps were done using Scikit-Learn’s Preprocessing module.

For our baselines, we implemented gradient-boosted trees found in Light-GBM and XGBoost. We used their respective Python libraries.

##### LGBM

We used LGBM(*Light gradient boosting Machine)* as our first classifier due to the effectiveness of this model in classification tasks in computational biology such as [39], [40], [41]. We determined the hyperparameters empirically. We trained the model for 1000 boosting rounds and used multiclass objectives since we have three classes. We used multiclass log loss as a loss function.

##### XGBoost

For our second model, we used XGBoost(*(eXtreme Gradient Boosting)* that has proven to be quite effective for various classification tasks in computational biology such as [42], [43], [44], [45]. We determined the hyperparameters empirically during training as well. We trained the model on the same GPU machine with 100 estimators and a max depth of 7.

##### ANN

We selected an ANN model for our third and main classification model. Such deep-learning models have been quite effective for various classification tasks in bioinformatics over the years such as [46], [47], [48], [49].

We implemented our ANN with the tools available in Pytorch. We used Adam as our optimizer with learning a rate of 1 *×* 10^*−*5^, amsgrad enabled, and weight decay of 0.001. We used Categorical Cross Entropy as our loss function with label smoothing of 0.1. We trained PRIEST for 100 epochs with batch size 32.

#### 7.1.5 Results

Our primary metrics of concern for classification are Accuracy, F1-Score, and ROC-AUC Score. From Table 9 that our ANN classifier model matches or outperforms other baseline models in terms of all scores including our metrics of concern. This is due to the availability of a large quantity of data.

**Table 9:** Classifier Performance on Test Set.

| Model | Accuracy | Precision | Recall | F1-Score | ROC-AUC Score |
| --- | --- | --- | --- | --- | --- |
| LGBM | 0.95 | 0.95 | 0.94 | 0.95 | 0.99 |
| XGBoost | **0.97** | 0.96 | 0.96 | 0.96 | **1.00** |
| ANN | **0.97** | **0.97** | **0.97** | **0.97** | **1.00** |

### 7.2 Mutation Prediction

#### 7.2.1 Performance Comparison of PRIEST and Other Baseline Models on Additional Metrics

The performance for other metrics such as Precision, Recall, AUROC, and AP of all the models used in our experiments is reported in Table 10. PRIEST has a higher Precision value than other baseline models for *k* **∈**{ 3, 6} . Additionally, for *k* = 9, it still has comparable performance, coming in second. For the AUROC score, PRIEST outperforms for all values of *k* compared to other baseline models. As For the AP score, we see a similar case to that of the Precision score where it has a higher AP value for *k* **∈**{3, 6 }. For Recall value, PRIEST performs competitively with other baseline models for all *k*.

**Table 10:**
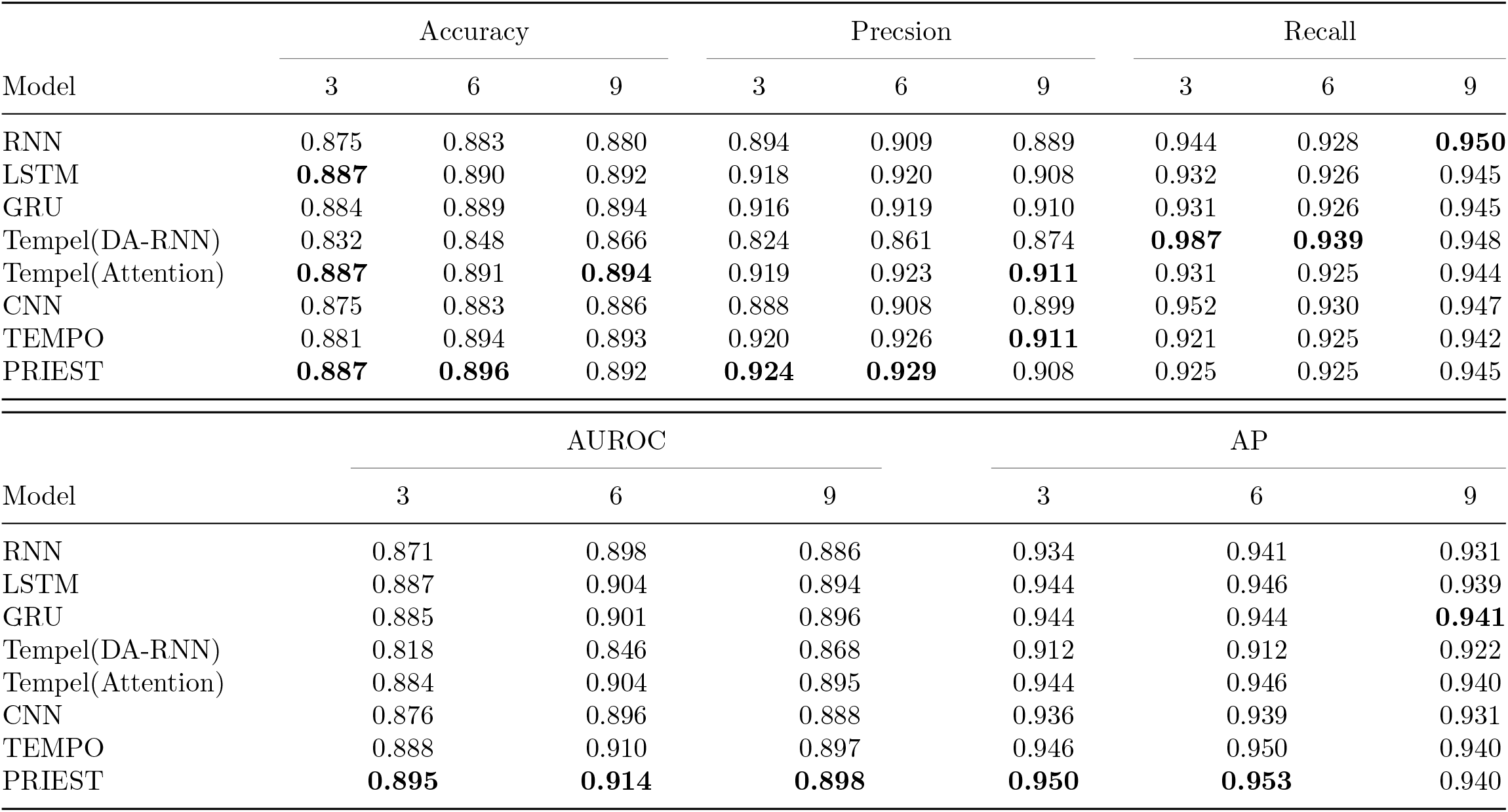
A comparison of evaluation metrics (Accuracy, Precision, Recall, AUROC, and AP) obtained by PRIEST and other benchmark models on independent test set.

**Table 11:** Pairwise comparison of Euclidean distances and cosine dissimilarity between clusters.

(a) Euclidean Distance
| Generated<br>Sequences<br>Actual<br>Sequences | Cluster 1 | Cluster 2 | Cluster 3 | Cluster 4 | Cluster 5 | Cluster 6 | Cluster 7 |
| --- | --- | --- | --- | --- | --- | --- | --- |
|  | Cluster 1 | Cluster 2 | Cluster 3 | Cluster 4 | Cluster 5 | Cluster 6 | Cluster 7 |
| Cluster 1 | **0.199** | 0.873 | 0.720 | - | 0.324 | - | 1.610 |
| Cluster 2 | 0.860 | **0.422** | 0.771 | - | 0.658 | - | 1.008 |
| Cluster 3 | 0.833 | 1.077 | 0.731 | - | 1.289 | - | 1.579 |
| Cluster 4 | 0.203 | 0.877 | 0.719 | - | 0.779 | - | 1.611 |
| Cluster 5 | 0.391 | 0.612 | **0.631** | - | **0.251** | - | 1.364 |
| Cluster 6 | 2.760 | 2.737 | 2.618 | - | 2.715 | - | 2.915 |
| Cluster 7 | 1.462 | 0.794 | 1.226 | - | 1.261 | - | **0.716** |

(b) Cosine Dissimilarity
| Generated<br>Sequences<br>Actual<br>Sequences | Cluster 1 | Cluster 2 | Cluster 3 | Cluster 4 | Cluster 5 | Cluster 6 | Cluster 7 |
| --- | --- | --- | --- | --- | --- | --- | --- |
|  | Cluster 1 | Cluster 2 | Cluster 3 | Cluster 4 | Cluster 5 | Cluster 6 | Cluster 7 |
| Cluster 1 | **0.006** | 0.131 | 0.084 | - | 0.016 | - | 0.397 |
| Cluster 2 | 0.128 | **0.035** | 0.124 | - | 0.082 | - | 0.162 |
| Cluster 3 | 0.119 | 0.224 | 0.106 | - | 0.268 | - | 0.406 |
| Cluster 4 | 0.007 | 0.131 | 0.083 | - | 0.016 | - | 0.396 |
| Cluster 5 | 0.024 | 0.073 | **0.079** | - | **0.012** | - | 0.302 |
| Cluster 6 | 0.846 | 0.886 | 0.832 | - | 0.188 | - | 0.913 |
| Cluster 7 | 0.336 | 1.001 | 0.259 | - | 0.261 | - | **0.076** |

### 7.3 Mutation Quality Testing

#### 7.3.1 Details of PCA parameter selection

#### 7.3.2 Details of GMM parameter selection

From graph 8a of BIC score vs. numbers of components of different covariance types, we observe that plot of BIC score of diagonal type covariance is lower than all other covariance types for components 2, 5, 6, 7, 8, and 9. So we selected this covariance type among all others. From graph 8b of Gradients of BIC score vs. the number of components, we observe that the gradient of spherical and full type of covariance changed after seven components. In comparison, the gradient of the diagonal and full type of covariance remained static after seven components. So, we finally fixed the number of components for the diagonal type of covariance to 7. Thus we finalized the covariance type to be diagonal and used 7 clusters or components for our GMM for testing the quality of generated mutated sequences by PRIEST.

